# Post-Learning Reactivation of Memories for Naturalistic Continuous Experiences

**DOI:** 10.64898/2026.10.20.760554

**Authors:** Nora Wolf, James W. Antony, Angelique I. Delarazan, Zachariah M. Reagh, Sebastian Michelmann, Charan Ranganath, Xiaonan L. Liu

## Abstract

Memories are not static records of the past; rather, the act of remembering can change how an event is recalled. Intriguingly, several studies also suggest that memories may be reactivated during rest. However, reactivation has largely been studied for spatial tasks and discrete stimuli, and we know little about whether spontaneous reactivation occurs for long timescale naturalistic events and how reactivation affects retention. Using fMRI, we show that neural activity patterns in the hippocampus and the Default Mode Network during encoding of narrative events were spontaneously reactivated during rest. Contrary to predictions, reactivation did not strengthen recall of movie details, and instead exploratory analyses suggested that reactivation in the hippocampus was associated with more gist-like recall one week later. These findings support the idea that memories for complex events may be spontaneously reactivated, but they suggest that rather than simply strengthening episodic memories, reactivation might help transform them.

**Significance statement:** We tend to think of memory as a process of recording and later playing back those experiences during retrieval, but some prior evidence suggests the brain may use downtime to process memories to help consolidate them. This paper uses functional MRI to show that the brain activation patterns present while people view films are spontaneously reactivated afterwards during rest. While there was no evidence to suggest that this reactivation strengthened memories, reactivation may be associated with more gist-like memory one week later. These findings suggest that the resting brain processes memories for complex events, but the results call for theories that account for transformations of memories beyond strengthening.

## Introduction

Many theories of memory consolidation propose that memories can be reactivated during offline states, transforming them and acting as a mechanism to save memories that might otherwise be forgotten (Rasch and Born, 2007; Tambini and Davachi, 2019; Mekbib and McDonough, 2025). Consistent with these theories, numerous studies have documented post-learning reactivation of sequences of hippocampal place cells in rodents during offline states (Carr et al., 2011; Foster, 2017), and some evidence in humans suggests that hippocampal representations of recent experiences may be spontaneously reactivated during sleep (Paller, 2024) and waking rest (Wamsley, 2022).

At present, however, it is unclear whether or how reactivation of place cells or representations of discrete stimuli translates to human memories for naturalistic events. Naturalistic events tend to have a schematic structure (e.g., parties, soccer games, etc.) and extend across long timescales. Evidence suggests that place cell sequence replays consist of fragmented, short segments of spatial trajectories (Davidson et al., 2009; Berners-Lee et al., 2022; Eliav et al., 2025), and studies of post-learning reactivation in humans generally focus on memory for arbitrary, unstructured associations (Tambini and Davachi, 2019; Mekbib and McDonough, 2025).

A second key question is the extent to which reactivation reliably enhances human episodic memory. Although targeted reactivation of learned associations during sleep can improve retention (Paller, 2017), the consequences of spontaneous reactivation of memories during post-learning rest appear to be more variable. Functional magnetic resonance imaging (fMRI) studies using multi-voxel pattern analysis have shown that spontaneous reactivation of learning-related activity patterns can predict better subsequent memory retention (Deuker et al., 2013; Staresina et al., 2013; Gruber et al., 2016; Yu et al., 2024), but reactivation has also been associated with forgetting (Tanrıverdi et al., 2023). Alternatively, some theories suggest that post-learning reactivation might transform memories, emphasizing shared content, rather than detailed information (e.g., Winocur et al., 2010; Lewis and Durrant, 2011; Moscovitch et al., 2016).

However, as noted above, there is limited empirical work on reactivation of memories during rest that unfold across a timescale that is representative of real-world episodic memories. Here, we used fMRI to investigate whether neural representations of a continuous, narrative experience are reactivated in the hippocampus and neocortex during periods of post-learning awake rest, and the effect of this reactivation on episodic memory. Our design (Figure 1) included both a pre-encoding rest period (acting as a baseline) and post-encoding rest period in order to unambiguously identify post-learning memory reactivation and to disentangle it from unspecific neural correlates of rest. Given the documented role of the default mode network (DMN) in episodic memory functions during both encoding and retrieval (Ranganath and Ritchey, 2012), we expected reactivation in the subnetworks of the DMN, as well as the hippocampus (cf. Kaplan et al., 2016).

**Figure 1.**
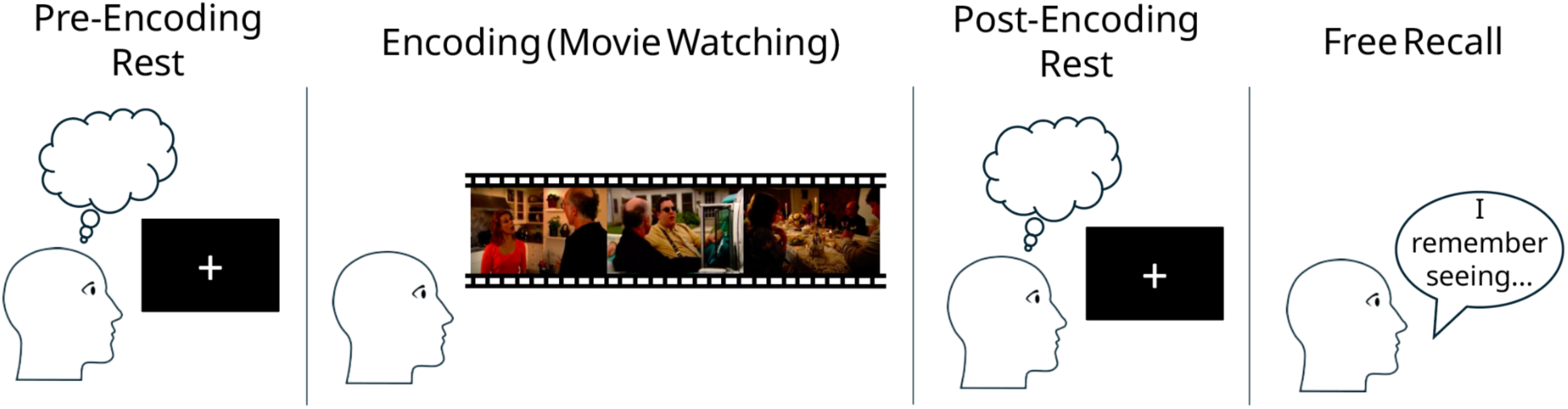
Experimental design. Participants completed a multi-phase fMRI experiment consisting of alternating rest and task periods during Session 1. Not shown here, participants returned one week later for a Session 2, during which they watched another episode from the same TV show and then completed an additional free recall of the original episode they had watched the week before.

In brief, participants completed a multi-phase fMRI experiment consisting of alternating rest and task periods. Participants first underwent an initial resting-state scan (Pre-Encoding Rest), which served as a control period with no recently encoded video content to reactivate. Participants then watched an episode of the TV show, *Curb Your Enthusiasm* (Encoding), while undergoing fMRI. Immediately afterward, they completed a second resting-state scan (Post-Encoding Rest), during which we expected neural reactivation of the recently encoded video stimulus. Next, participants freely recalled what they remembered from the episode while inside the scanner. One week later, participants returned for a second session, during which they watched and recalled another episode from the same TV show and then again freely recalled the original episode they had watched during Session 1.

## Methods

### Participants

Participants were recruited from the University of California, Davis community and the surrounding area. The study was part of a larger study looking at age-related differences in episodic memory (see Bouffard et al., 2026; Delarazan et al., 2026); in this paper, we only analyzed the subset of neural data from participants under 40 years old. Exclusion criteria included magnetic bodily implants, claustrophobia, a history of major psychiatric or neurological disorders, a concussion in the past 6 months, untreated diabetes or hypertension, current drug or alcohol abuse, uncorrected hearing, and uncorrected vision. The experiment protocol and consent forms were approved by the Institutional Review Board of the University of California, Davis. Participants received $20/hour as compensation for their time.

Of the 39 young participants initially recruited, six participants were excluded because they had unusable functional data during at least one scan required for the analyses due to issues such as scanner malfunction causing image distortions, three participants were excluded because of excessive motion during at least one functional scan required for analyses, and one participant was excluded due to a file transfer problem, resulting in a final sample size of 29 participants (21 female, 8 male, age range 18-28 years). In addition, one participant was missing recall data for the second session and thus excluded from the memory analyses involving this session, and one participant recalled no detail units and was thus excluded from the gist-to-detail ratio analyses.

### Stimuli

Participants watched a 26-minute episode of a television “slice-of-life” sitcom (HBO’s *Curb Your Enthusiasm*, S01E07 titled “AAMCO”). The audio-visual stimulus chosen was selected to simulate everyday life experiences and be engaging to a wide range of ages.

### Procedure

Participants were briefed on the experiment and provided written informed consent. Participants then completed an MRI screening form, compensation form, and demographics questionnaires. Upon successful screening for the experiment, participants were escorted to the MRI facility and were prepared to undergo scanning.

High-resolution structural images and functional imaging runs were acquired (see *fMRI Data Acquisition*). As shown in Figure 1, participants first underwent an 8-minute long rest period scan (Pre-Encoding Rest), which was used as a control rest period for reactivation. Participants were then instructed to pay attention while watching an episode of *Curb Your Enthusiasm* titled “AAMCO,” and told that they would be recalling the episode in detail afterwards. After viewing the 26-minute TV episode (Encoding), participants underwent another 8-minute rest period scan (Post-Encoding Rest), which was used to identify reactivation of the just-seen video stimulus. Participants were then instructed to recall the episode in as much detail as possible while their audio was recorded (Free Recall). They were asked to recall details in the original viewing order when possible but were told remembering something out of order was acceptable. Participants then underwent a third 8-minute rest period scan, and finally watched an 11-minute clip from the start of the movie *Catch Me If You Can*. During the resting period scans, participants were told to relax and to freely think about anything they would like.

Participants returned a week later, either to just complete behavioral tasks (n=9) or for an additional fMRI session (n=20). During this session, participants completed some personality ratings of the characters from the “AAMCO” episode before watching another episode from *Curb Your Enthusiasm* named “The Wire.” They then freely recalled the episode “The Wire,” and completed additional personality ratings of the characters from *Curb Your Enthusiasm*. Finally, participants were asked to again freely recall what they remembered from the “AAMCO” episode that they had watched the previous week. Of the tasks completed during this Session 2, only the free recall was used in the analyses reported below.

### Event boundary time point data collection and preparation

In order to determine event boundaries in the main video stimulus, a separate group of 20 participants watched the “AAMCO” episode and then segmented the video by indicating every time, in their opinion, one meaningful event had ended and another had begun (Newtson, 1973; Zacks and Swallow, 2007).

The ratio of participants who perceived an event boundary within the same three-second integration window was computed for each time point in the video. Peaks in the time course with >=50% agreement across participants, and a temporal distance of at least six seconds, were marked as event boundaries; if two peaks occurred within this six-second window, only the higher peak was selected. In addition, the time point at which the opening credits ended and the narrative began was marked as an event boundary.

### fMRI acquisition

MRI scanning for the primary dataset was performed on a 3T Siemens Skyra scanner with a 32-channel head coil. T1-weighted structural images were acquired using a magnetization prepared rapid acquisition gradient echo (MPRAGE) pulse sequence (TR = 2100 ms; TE = 2.98 ms; flip angle = 7°; field of view = 256 mm^2^; image matrix = 256 x 256, 208 sagittal slices, voxel size = 1 mm^3^). Functional images were acquired using a T2*-weighted gradient EPI sequence (TR = 1,220 ms; TE = 24 ms; field of view = 192 mm^2^ ; image matrix = 64 x 64; flip angle = 67°; multiband factor = 2; 38 interleaved axial slices, voxel size = 3 mm^3^ isotropic, P » A phase encoding). A functional scan of reverse phase encoding polarity (A » P) was acquired to correct for phase encoding distortion.

### fMRI processing

Preprocessing was performed using the standardized fMRIPrep pipeline. Specifically, MRI data were preprocessed using fMRIPrep 23.1.2 (Esteban et al., 2019, RRID:SCR_016216), which is based on Nipype 1.8.6 (Gorgolewski et al., 2011, RRID:SCR_002502). Many internal operations of fMRIPrep use Nilearn 0.10.1 (Abraham et al., 2014, RRID:SCR_001362). Each T1-weighted (T1w) image was corrected for intensity non-uniformity (INU) with N4BiasFieldCorrection (Tustison et al., 2010), distributed with ANTs (Avants et al., 2008, RRID:SCR_004757). The T1w-reference was then skull-stripped with a Nipype implementation of the antsBrainExtraction.sh workflow (from ANTs), using OASIS30ANTs as target template. Brain tissue segmentation of cerebrospinal fluid (CSF), white-matter (WM) and gray-matter (GM) was performed on the brain-extracted T1w using fast (FSL, RRID:SCR_002823, Zhang et al., 2001). An anatomical T1w-reference map was computed after registration of 2 T1w images (after INU-correction) using mri_robust_template (FreeSurfer 7.3.2, Reuter et al., 2010). Brain surfaces were reconstructed using recon-all (FreeSurfer 7.3.2, RRID:SCR_001847, Dale et al., 1999), and the brain mask estimated previously was refined with a custom variation of the method to reconcile ANTs-derived and FreeSurfer-derived segmentations of the cortical gray-matter of Mindboggle (RRID:SCR_002438, Klein et al., 2017). Volume-based spatial normalization to one standard space (MNI152NLin2009cAsym) was performed through nonlinear registration with antsRegistration (from ANTs), using brain-extracted versions of both T1w reference and the T1w template.

For each of the BOLD runs per subject (across all tasks and sessions), first, a reference volume and its skull-stripped version were generated using a custom methodology of fMRIPrep. Head-motion parameters with respect to the BOLD reference (transformation matrices, and six corresponding rotation and translation parameters) were estimated before any spatiotemporal filtering using mcflirt (FSL, Jenkinson et al., 2002). A B0-nonuniformity map (fieldmap) was estimated based on two echo-planar imaging (EPI) references with topup (Andersson et al., 2003, FSL). The estimated fieldmap was then aligned with rigid-registration to the target EPI (echo-planar imaging) reference run. The field coefficients were mapped on to the reference EPI using the transform. BOLD runs were slice-time corrected to 0.569s (0.5 of slice acquisition range 0s-1.14s) using 3dTshift from AFNI (Cox and Hyde, 1997, RRID:SCR_005927). The BOLD reference was then co-registered to the T1w reference using bbregister (FreeSurfer) which implements boundary-based registration (Greve and Fischl, 2009). Co-registration was configured with six degrees of freedom. Several confounding time series were calculated based on the preprocessed BOLD: framewise displacement (FD), DVARS and three region-wise global signals. FD was computed using two formulations following Power (absolute sum of relative motions (Power et al., 2014)) and Jenkinson (relative root mean square displacement between affines (Jenkinson et al., 2002)). FD and DVARS were calculated for each functional run, both using their implementations in Nipype (following the definitions by Power et al., 2014). The three global signals were extracted within the CSF, the WM, and the whole-brain masks. The BOLD time series were resampled into standard space, generating a preprocessed BOLD run in MNI152NLin2009cAsym space. All resampling was performed in a single interpolation step by composing all pertinent transformations (i.e., head-motion transform matrices, susceptibility distortion correction, and co-registrations to anatomical and output spaces). Gridded (volumetric) resampling was performed using antsApplyTransforms (ANTs), configured with Lanczos interpolation to minimize the smoothing effects of other kernels (Lanczos, 1964). Non-gridded (surface) resampling was performed using mri_vol2surf (FreeSurfer).

Following fMRIPrep preprocessing, an across-run averaged subject mask was applied to each participant’s data, and data for each run were re-sampled to a 2-mm isotropic resolution. Confound variables (six rotation and translation confounds and FD) were regressed out from the fMRI time course, and each voxel’s time series was *z*-scored across time. Time points with >0.5-mm FD or spikes >3 standard deviations from the mean of the global time series were removed from analyses. All calculations were performed in volume space. Event boundary time points from the video used in analyses were shifted by six TRs (7.32 s) to align with the neural time series due to hemodynamic delay.

### Regions of Interest (ROIs)

Normalized time series were parcellated using an extended HCP-MMP atlas, which contained both cortical and subcortical parcels in volumetric space (Huang et al., 2021, 2022). Parcel-level network identity was then assigned using the cortico-hippocampal network classification derived from data-driven resting-state functional connectivity analyses (Barnett et al., 2021). Our *a priori* regions of interest (ROIs) were the hippocampus and the Default Mode Network (DMN) and its three subnetworks: the Posterior Medial Network (PMN), which includes the posterior cingulate cortex, anterior retrosplenial cortex, posterior angular gyrus, and dorsal prefrontal cortex, the Anterior Temporal Network (ATN), which includes the temporopolar cortex, lateral orbitofrontal cortex, temporoparietal junction, and dorsomedial prefrontal cortex, and the Medial Prefrontal Network (MPN), which includes medial prefrontal cortex regions and the entorhinal cortex (Barnett et al., 2021). In addition, the hippocampal head, body and tail were defined from probabilistic maps created by Ritchey et al. (2015).

### Quantifying reactivation

To quantify reactivation for both the Post-Encoding Reactivation (correlating Encoding with Post-Encoding Rest) and Pre-Encoding Control (correlating Encoding with Pre-Encoding Rest) conditions, we computed time-by-time pattern similarity matrices between resting-state and encoding multivoxel activity patterns for each participant and ROI as follows:

We first extracted voxel activity patterns within each parcel in a given ROI for every resting time point and every encoding time point. We then computed the Pearson correlation coefficient between the neural activity pattern at each resting time point and the neural activity pattern at each encoding time point. This produced a time-by-time correlation matrix that captured how strongly every encoding point was reactivated at every resting time point. We averaged these matrices across all parcels within an ROI to yield a single correlation matrix per ROI. Assuming a single encoding time point was being reactivated at any given moment during rest, we selected the highest pattern similarity value at each resting-state time point from the averaged time-by-time correlation matrix as the most likely candidate for reactivation during that resting-state time point.

Alternatively, rather than a single encoding time point being reactivated at any given moment during rest, it is possible that multiple time points were reactivated simultaneously. Thus, we repeated the analysis on all positive correlations in the matrix, instead of just the highest one at each resting time point.

Finally, we Fisher *z*-transformed these selected correlations and computed their mean to generate a single metric of mean reactivation strength per participant for each ROI and condition.

### Quantifying reactivation of events

We ran a third variation of the analysis in which we looked for reactivation of events, rather than individual time points from encoding. To quantify reactivation of events from the encoding experience, rather than each individual time point, we followed the same procedure as described above but instead broke the neural activity patterns of each encoding run into event-averaged segments. After extracting voxel activity patterns for each encoding time point, we segmented the encoding time series, based on the time points marking the beginning and end of events that had been identified by a separate group of participants (see *Event boundary time point data collection and preparation*). We then averaged the neural activity patterns across time points within each event, resulting in one pattern of brain activity for each event.

We then correlated the neural activity patterns at each resting time point with the neural activity patterns of each event-averaged segment, rather than each individual encoding time point, to create correlation matrices. Selecting the highest correlation value at each resting time point now identified which event was most strongly reactivated at each resting time point, rather than which specific encoding time point was most strongly reactivated.

Finally, to compare reactivation of these events to reactivation of event boundary periods and mid-event periods, we repeated the reactivation strength calculation with modified averaging windows from the encoding run: namely, the boundary period (approximately 5 seconds before to approximately 5 seconds after the boundary), and a mid-event period used as an additional control (approximately 5 seconds before to approximately 5 seconds after the midpoint of the event), in addition to the entire event (from the event onset boundary to event offset boundary). Because these three encoding window types resulted in the same number of encoding segments, their reactivation strengths could be compared directly to each other.

### Relating reactivation to later recall

To score free recall, a manual scoring method was adapted from the Autobiographical Memory Interview (Levine et al., 2002). The “AAMCO” TV episode was broken into 195 units, and each unit was given a description. Each recall transcript was then scored for the number of units recalled, with a maximum possible score of 195.

Each transcript received both a score for the number of ‘gist units’ recalled and a score for the number of ‘detail units’ recalled, with gist units being awarded when a participant conveyed the general meaning of a unit description, and detail units being awarded when the participant additionally recalled the specific components (who/where/what). Any detail unit awarded necessarily implied the corresponding gist unit was also awarded, whereas gist units could be awarded in the absence of detail. The gist-to-detail ratio was calculated by dividing the number of gist units a participant recalled by the number of detail units they recalled.

Due to the possibility that participants might have been uncomfortable talking about the last scene due to its content, we excluded it from memory analyses for a more accurate measure of memory, though including memory scores from the last scene did not materially change any results reported.

To assess the relationship between reactivation and memory, we correlated the mean level of reactivation for each participant with these memory measures. To account for any subject-level differences in baseline neural similarity, we calculated mean reactivation strength for each participant as the difference between their post-encoding reactivation mean and their pre-encoding control mean (described above). These subject-level reactivation values (corresponding to those plotted in Figure 3 for time point-based analyses and Figure 4 for event-based analyses) in the hippocampus and DMN were then paired with each participant’s memory scores (gist units, detail units, and gist-to-detail ratio) across Session 1 and Session 2 free recall tasks.

### Statistical analysis

All reactivation group analyses were performed on Fisher *z*-transformed values, and all reactivation data are reported as Fisher *z*-transformation values. To test our directional hypothesis of whether Post-Encoding Reactivation was significantly higher than Pre-Encoding Control, we performed one-sided paired t-tests.

To control for multiple comparisons across the three different ways we tested for reactivation (selecting the highest encoding time point correlation at each resting time point, selecting all positive encoding time point correlations, and selecting the highest event template correlation at each resting time point), each tested in both the hippocampus and DMN, for a total of 6 comparisons, we applied False Discovery Rate (FDR) correction using the Benjamini-Hochberg procedure with a significance threshold of *Q* < 0.05. Subregion/subnetwork tests were subsequently conducted uncorrected (*α* < 0.05), contingent upon significant FDR-corrected main effects.

To compare reactivation strength between entire events, event boundary periods, and mid-event periods, we conducted a one-way repeated-measures ANOVA across these three window conditions.

Finally, to correlate reactivation strength with memory, we conducted Pearson’s correlation coefficients (*r*). FDR correction was applied across all memory performance comparisons (number of gist units recalled and number of detail units recalled, for both Session 1 and Session 2, for reactivation in both the hippocampus and the DMN), with a significance threshold of *Q* < 0.05, but no *a priori* hypotheses reached significance. We subsequently conducted uncorrected exploratory analyses for correlations involving the gist-to-detail ratio.

## Results

### Evidence and quantification of reactivation

Based on research suggesting that hippocampal representations of recent experience are reactivated during waking rest (Foster, 2017; Tambini and Davachi, 2019), we hypothesized that neural activity patterns observed during movie watching would be reactivated during post-learning rest. We tested this by investigating whether there was a higher degree of similarity between the neural activity patterns during encoding and a post-encoding resting state scan (Post-Encoding Reactivation) than between encoding and a resting state scan taken before participants had seen the movie (Pre-Encoding Control) (Figure 2). Findings suggest that neural activity during encoding can be similar to a pre-encoding period (Sadeh et al., 2019), which may lead to smaller differences between the two conditions; nevertheless, we expect the similarity to be greater for the post-encoding period than for the pre-encoding period.

**Figure 2.**
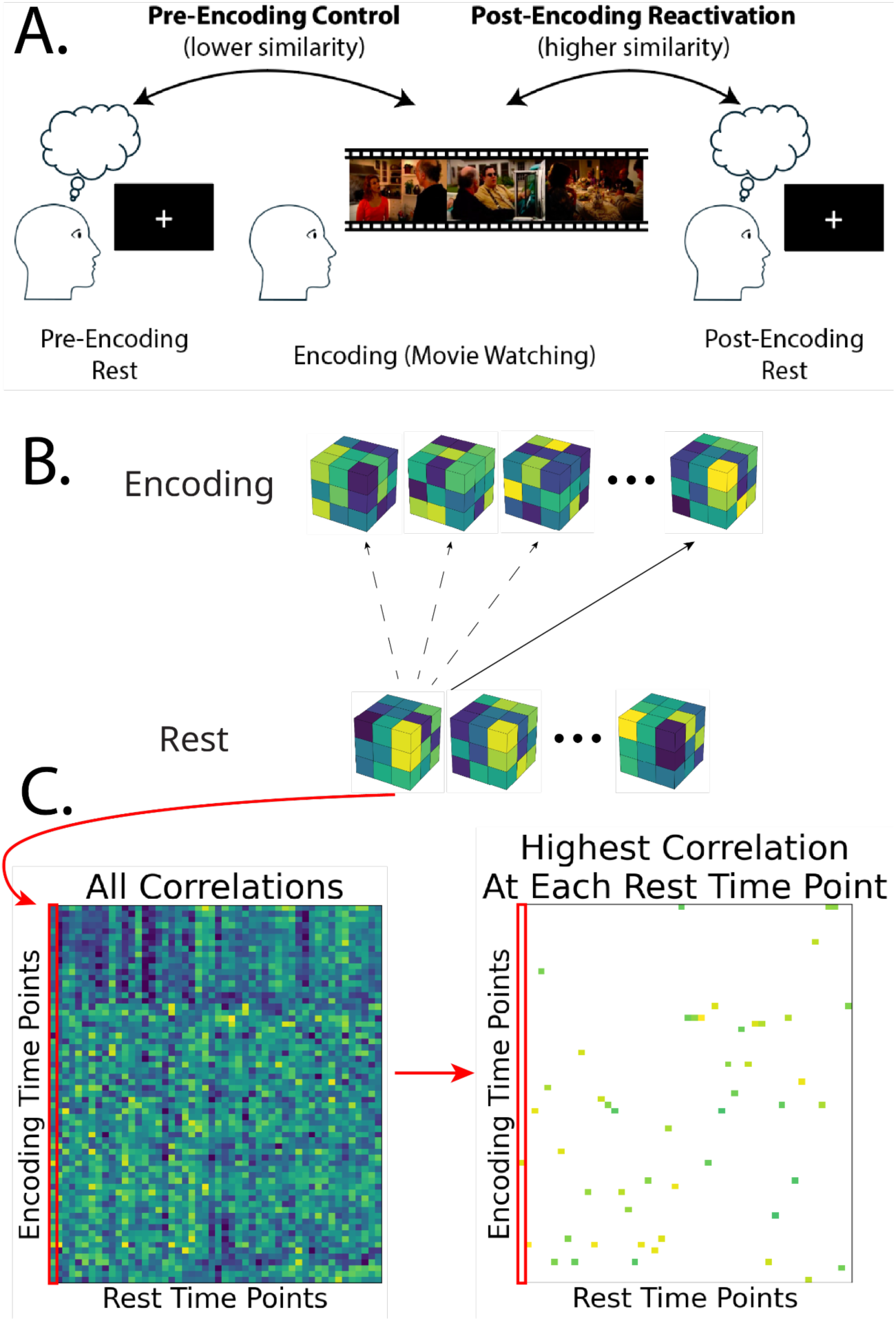
Reactivation analysis. A. If reactivation of the encoding experience occurred during post-encoding rest, we would expect a higher degree of similarity in neural activity between the post-encoding rest period and the encoding period (Post-Encoding Reactivation comparison) than between the pre-encoding rest period and the encoding period (Pre-Encoding Control comparison). B. To quantify the level of similarity between the encoding experience and a rest period, we correlated the neural activity patterns at every time point during the resting state scans with every neural activity pattern recorded during movie encoding. Next, for each resting time point, we selected the encoding time point with the highest correlation (i.e., the most similar neural activity pattern) as the candidate time point for reactivation. C. Example of resulting correlation matrix with all resting time point and encoding time point correlations, and subsequent correlation matrix with highest correlation at each resting time point selected.

We assumed that any neural pattern observed during movie watching could be reactivated at any time point during post-encoding rest. To quantify reactivation, we first extracted voxel activity patterns at every resting time point and at every time point during encoding. Next, for each resting time point, we calculated neural pattern similarity (using the Pearson’s correlation coefficient) with each of the activity patterns observed during encoding. The encoding time point with the highest pattern similarity value was selected as the most likely candidate for reactivation during that resting-state time point.

We would not expect reactivation to occur at every resting time point. However, if post-encoding reactivation occurred at some time points, then we should expect the mean of the selected highest correlation values during post-encoding rest to be higher than the mean of the selected highest correlation values in the pre-encoding rest control.

Consistent with our directional hypothesis, one-tailed t-tests revealed significantly higher correlations for the post-encoding reactivation condition than for the pre-encoding control in both the hippocampus, *t*(28) = 2.08, *p* = .023, *p*_adj_ = .035, and DMN, *t*(28) = 2.60, *p* = .007, *p*_adj_ = .016 (Figure 3). Following up on these significant whole-structure effects, uncorrected post-hoc analyses were conducted to examine anatomical specificity across subregions and subnetworks. Looking at the head, body and tail of the hippocampus separately, the post-encoding reactivation condition also showed significantly higher correlations than the control condition in the hippocampal head, *t*(28) = 2.05, *p* = .025, but not in the hippocampal body, *t*(28) = 1.54, *p* = .067, or tail, *t*(28) = 1.46, *p* = .078. Looking at the subnetworks of the DMN, post-encoding reactivation showed significantly higher correlations than the control in the PMN, *t*(28) = 3.02, *p* = .003, ATN, *t*(28) = 2.60, *p* = .007, and MPN, *t*(28) = 2.35, *p* = .013.

**Figure 3.**
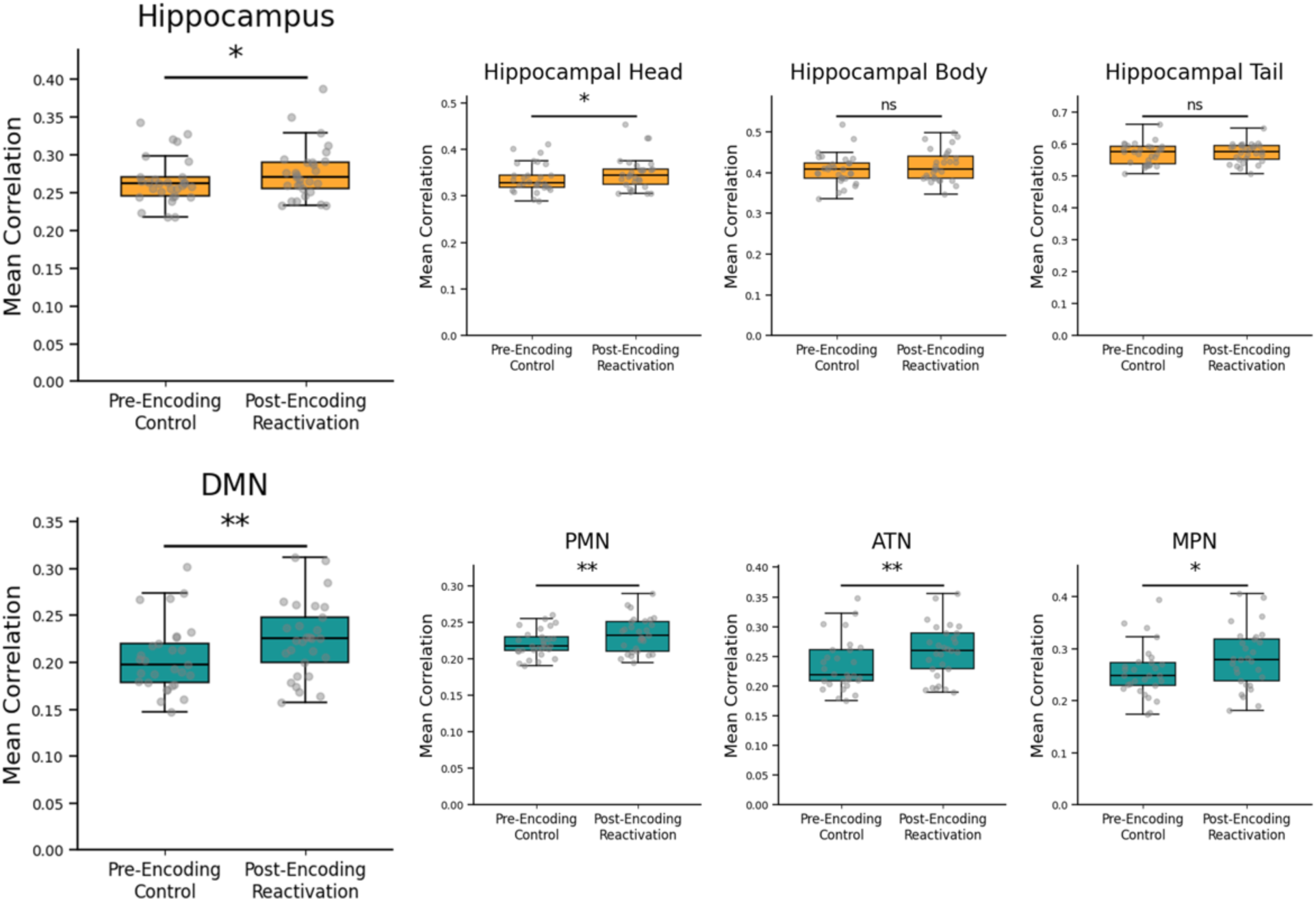
Reactivation of encoding time points during rest. Significantly higher correlations were found for post-encoding reactivation compared to the pre-encoding control condition in both the hippocampus and DMN. Each dot overlaid on the boxplot represents a participant. Asterisks denote statistical significance based on uncorrected *p*-values (* *p* < 0.05, ** *p* < 0.01, ns = not significant).

Rather than a single encoding time point being reactivated at any given moment during rest, it is possible that multiple time points were reactivated simultaneously. Thus, we repeated the analysis on all positive correlations, instead of just the highest one at each resting time point. This analysis additionally provides a more complete comparison between the post-encoding reactivation and pre-encoding control conditions, incorporating a much larger number of data points rather than selecting just the strongest correlations. We found significant levels of reactivation in the DMN, *t*(28) = 2.57, *p* = .008, *p*_adj_ = .016, but the difference between post-encoding reactivation and the pre-encoding control did not reach significance in the hippocampus, *t*(28) = 1.65, *p* = .055.

### Evidence and quantification of reactivation of events and event boundaries

Prior research suggests that rather than encoding every moment in time, we segment continuous experiences into events that become consolidated into long-term memory (Baldassano et al., 2017; Zacks, 2020). Because we hypothesized that reactivation for episodic memories may happen at a slower timescale, and studies have shown that neural activity patterns averaged across the time course of an event are reinstated during recall (Chen et al., 2017), we next examined whether we would find reactivation of events rather than individual time points.

We conducted the same analysis as described above comparing correlations in the Post-Encoding Reactivation condition to the Pre-Encoding Control condition, but tested for the reactivation of each mean event pattern rather than each encoding time point. Since neural activity has been shown to be relatively stable within an event, we averaged the neural activity patterns across each event to get an ‘event template’ of the typical activity pattern of that event, using time points identified by a separate group of participants for the start and end of events. We then correlated the neural activity patterns of each resting time point with each of these event-averaged ‘templates,’ and selected the highest correlation at each resting time point to get a candidate for which event was most likely reactivated at that resting time point. As before, we expected the mean selected correlations to be higher for the Post-Encoding Reactivation comparison than the Pre-Encoding Control comparison.

As shown in Figure 4, in the hippocampus, post-encoding reactivation correlations were significantly higher than for the pre-encoding control, *t*(28) = 1.97, *p* = .029, *p*_adj_ = .035, and uncorrected post-hoc analyses to decompose this main effect additionally revealed significant reactivation in the hippocampal head, *t*(28) = 2.20, *p* = .018, but not in the hippocampal body, *t*(28) = 1.02, *p* = .158, or tail, *t*(28) = 1.33, *p* = .098. In the DMN, the post-encoding reactivation condition was significantly higher than for the pre-encoding control, *t*(28) = 2.65, *p* = .006, *p*_adj_ = .016, and uncorrected post-hoc analyses revealed significant reactivation in all subnetworks of the DMN (PMN, *t*(28) = 2.19, *p* = .018, ATN, *t*(28) = 2.58, *p* = .008, and MPN, *t*(28) = 2.57, *p* = .008).

**Figure 4.**
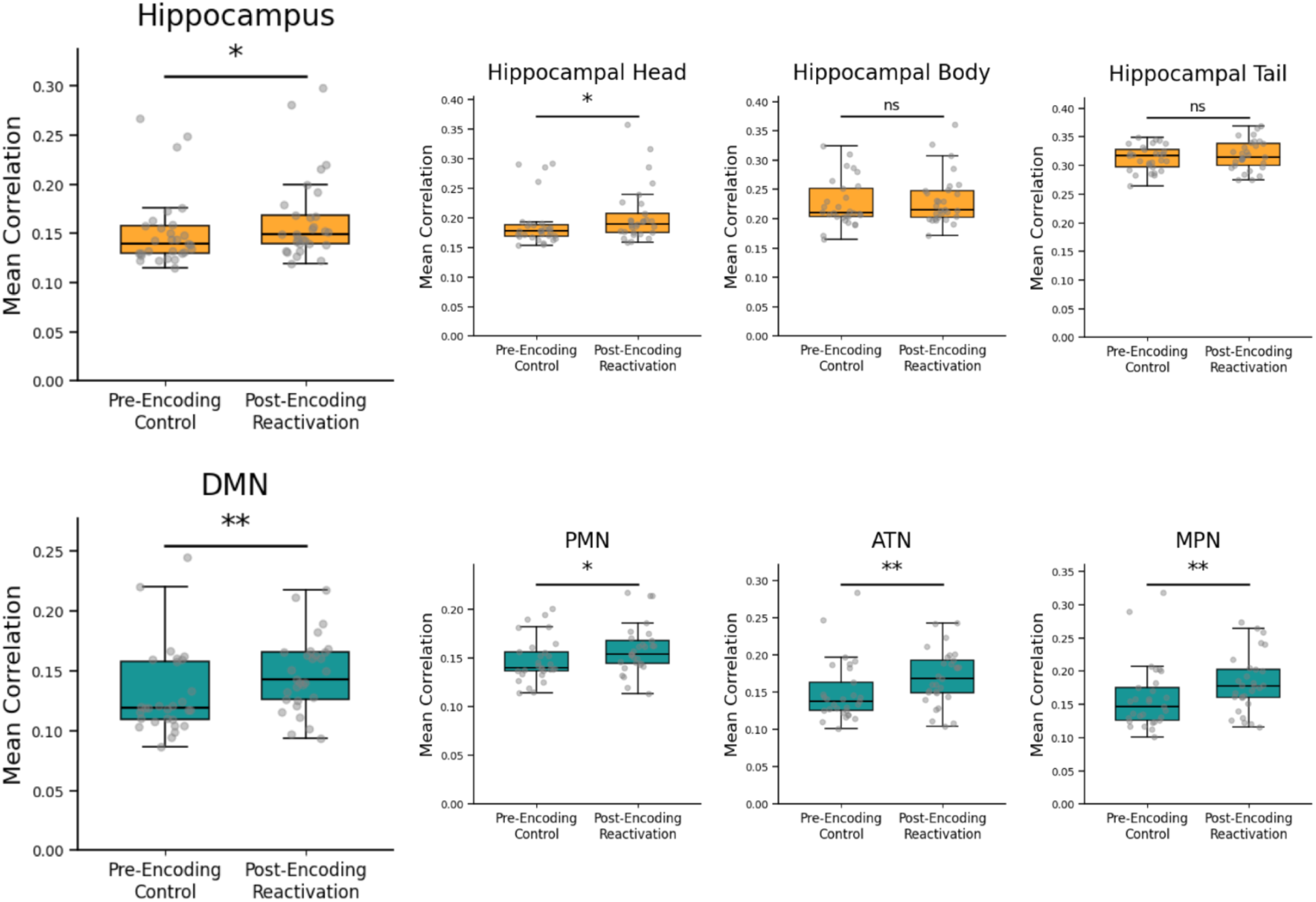
Reactivation of encoding event-averaged templates during rest. Significantly higher correlations were found for post-encoding reactivation compared to the pre-encoding control condition in both the hippocampus and DMN. Each dot overlaid on the boxplot represents a participant. Asterisks denote statistical significance based on uncorrected *p*-values (* *p* < 0.05, ** *p* < 0.01, ns = not significant).

Alternatively, it is possible that because event representations are encoded at event boundaries, the periods at which changes in context occur and one event begins and another ends (Zacks, 2020), we would find stronger levels of reactivation of the neural activity patterns at event boundaries rather than of the events as a whole. To test this, we computed correlations for the Post-Encoding Reactivation condition using three different averaging windows from the encoding run: the boundary period (approximately 5 seconds before to approximately 5 seconds after the boundary), the entire event (from the event onset boundary to event offset boundary), and a mid-event period used as an additional control (approximately 5 seconds before to approximately 5 seconds after the midpoint of the event).

The boundary-averaged periods from encoding did not show significantly different levels of reactivation during the post-encoding rest period than event-averaged or mid-event-averaged periods in both in the hippocampus, *F*(2, 56) = 0.36, *p* = .700, and DMN, *F*(2, 56) = 1.23, *p* = .301.

### Relating reactivation to memory

Finally, to assess the behavioral relevance of reactivation, we asked whether reactivation strength was correlated with overall memory performance. To do this, we calculated each participant’s mean reactivation strength as the difference between their post-encoding reactivation mean and their pre-encoding control mean (calculated in the previous sections), and then correlated this value with the total number of idea units they recalled. We did this separately for both the number of gist units and the number of detail units recalled, looking at recall both during the same day (Session 1) and a week later (Session 2) in both the hippocampus and the DMN. We first report the results for the reactivation analysis on individual time points from the encoding experience (using the reactivation results shown in Figure 3).

In the hippocampus, the correlations for each of the four comparisons trended negatively but were not significant (all *r*’s ≥ -.16, all *p*’s ≥ .414). On the other hand, in the DMN, the correlations for each of the four comparisons was positive but also did not reach significance (all *r*’s ≤ .12, all *p*’s ≥ .532).

Having found no relationship between reactivation and memory performance, we conducted an exploratory analysis to test whether reactivation was instead associated with memory quality, namely, how gist-like the memories were. Since these exploratory analyses were not corrected for multiple comparisons, these findings require replication in a future study to confirm their validity. We correlated the mean reactivation strength of each participant with their gist-to-detail ratio (calculated as the ratio between the number of gist units recalled to the number of detail units recalled). As shown in Figure 5, mean reactivation strength correlated positively with the gist-to-detail ratio in both Session 1 and Session 2 in the hippocampus, with a significant (uncorrected) positive relationship between hippocampal reactivation strength and the gist-to-detail ratio in Session 2, *r* = .47, *p* = .013, but not Session 1, *r* = .30, *p* = .116. There was no significant relationship in the DMN, for both Session 1, *r* = .00, *p* = .998, and Session 2, *r* = .16, *p* = .435.

**Figure 5.**
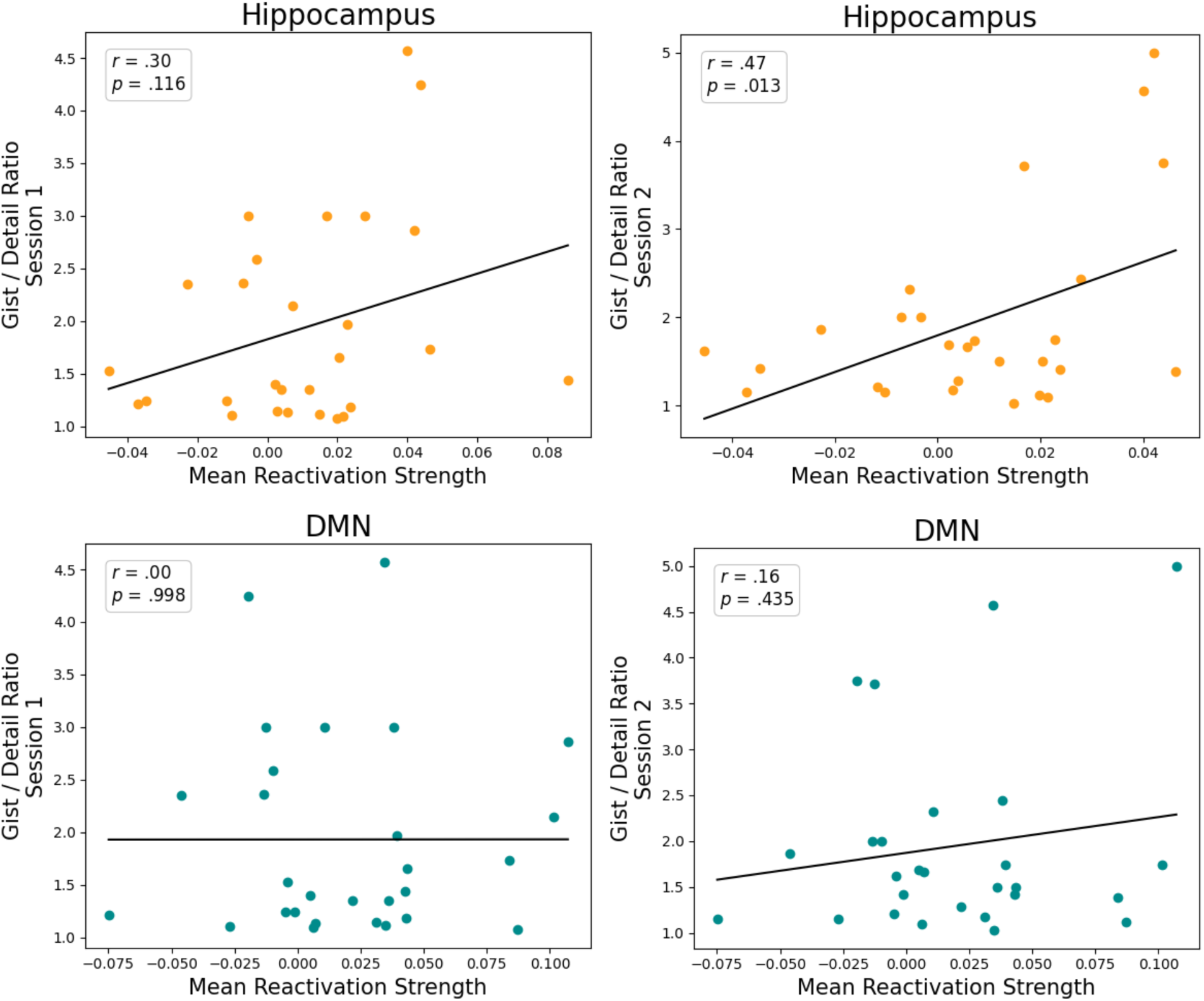
Correlations between mean reactivation strength of time points and gist-to-detail ratio for recalled units in Session 1 and Session 2, in the hippocampus and DMN.

Repeating these same analyses but for reactivation of event-averaged voxel patterns (shown in Figure 4), rather than time point-based patterns, we found similar results. There was no significant correlation between reactivation and memory performance in either the hippocampus or the DMN. Repeating the exploratory analyses on memory quality, we found the correlation between reactivation and the gist-to-detail ratio was positive for both the hippocampus and DMN in Session 1 and Session 2, with a significant (uncorrected) correlation between reactivation in the hippocampus and gist-like memory during Session 2, *r* = .43, *p* = .024.

## Discussion

We sought to extend findings on awake reactivation to the timescale of episodic memory. We found that multi-voxel patterns observed during encoding of a narrative film were reactivated during post-learning rest in both the hippocampus and cortical regions in the DMN. Critically, these correlations were higher than baseline correlations observed between movie-watching and pre-encoding resting-state activity. In addition to reactivation of discrete time points, we observed reactivation of event-averaged activity patterns in the hippocampus and DMN. Although post-learning reactivation was not correlated with subsequent memory performance, exploratory analyses suggested that reactivation in the hippocampus may be associated with more gist-like memory one week later. These results provide evidence that brain activity patterns in cortico-hippocampal networks observed during naturalistic experiences are spontaneously reactivated during waking rest, and that reactivation could play a role in memory transformation, rather than simply strengthening or weakening memories.

Sequential replay of place cells in the rodent hippocampus, often coinciding with sharp wave ripples, has been thought to facilitate hippocampal-cortical interactions that support memory consolidation (Kaplan et al., 2016; Ólafsdóttir et al., 2018). In humans, a number of fMRI studies have examined how hippocampal and neocortical activity during encoding of items or simple associations relates to activity during post-learning rest. Post-learning changes in functional connectivity between the hippocampus and neocortical areas that were engaged during learning are reliably correlated with subsequent memory performance (Tambini et al., 2010; Schlichting and Preston, 2014; Gruber et al., 2016; Collins and Dickerson, 2019; Folvik et al., 2023), and reactivation has been associated with increased hippocampal connectivity with the DMN (Huang et al., 2024). Furthermore, multivariate pattern analyses have found reactivation in both the hippocampus (Gruber et al., 2016; Schapiro et al., 2018; Schuck and Niv, 2019; Hindy et al., 2026) and in cortical regions, including the visual cortex (Deuker et al., 2013; Wittkuhn and Schuck, 2021) and regions with strong interactions with the hippocampus such as the medial prefrontal cortex (Yu et al., 2024).

The present study examined post-learning reactivation of narrative events, which are more structured and have a much longer timescale than the typical items examined in prior fMRI studies. We found evidence that activity patterns of individual time points during encoding, as well as activity patterns averaged across entire events (extending up to several minutes), were transiently reactivated during rest in both the hippocampus and DMN. Our findings of reactivation in the DMN as well as in the hippocampus are consistent with the DMN’s role in episodic memory (Ranganath and Ritchey, 2012; Rugg and Vilberg, 2013) and its high degree of functional connectivity with the hippocampus (Kahn et al., 2008; Wang et al., 2016; Barnett et al., 2021; Reznik et al., 2023; Braga, 2025). Along the longitudinal extent of the hippocampus, reactivation effects were only significant in the anterior hippocampus. Additionally, we found no evidence for preferential reactivation of activity patterns around event boundaries, as opposed to whole-event or mid-event activity patterns. Previous studies have indicated that the DMN (Hasson et al., 2015; Chen et al., 2016; Zuo et al., 2020) and anterior hippocampus (Brunec et al., 2018; Bouffard et al., 2026) may preferentially encode information across relatively coarse spatiotemporal scales, and that the anterior hippocampus may preferentially encode the gist of events (Robin and Moscovitch, 2017). Collectively, these findings suggest that reactivation preserves the longer-timescale elements of episodes.

Post-learning reactivation of memories is often interpreted to support memory consolidation, and the simplest prediction to emerge from this account is that more reactivation should predict better memory performance. Whereas some studies in humans have found results consistent with this prediction (Deuker et al., 2013; Staresina et al., 2013; Schlichting and Preston, 2014; Gruber et al., 2016), others have found that reactivation can be associated with forgetting, or that it can selectively prioritize retention of weakly learned information. For instance, Tanrıverdi et al. (2023) showed that co-reactivations of the same stimulus in the hippocampus and fusiform face area were associated with worse recognition memory performance one week later and that these co-reactivations were specifically associated with trials that were initially correctly recognized immediately after encoding but then forgotten later on. In addition, Schapiro et al. (2018) found that items remembered less well were more likely to be reactivated during subsequent waking rest, suggesting that weakly encoded information gets prioritized for memory reactivation. However, they found marginal evidence that reactivation was associated with worse memory performance on a later test when participants did not sleep before being tested. This suggests that awake reactivation may not improve memory in the short term, with benefits emerging only after a longer delay that includes sleep. Finally, Zhang et al. (2018) identified reactivation using intracranial electrode recordings in humans, and found no difference in spontaneous awake reactivation between remembered and forgotten items.

In contrast to standard consolidation theories, memory transformation theories (Winocur et al., 2010; Winocur and Moscovitch, 2011; Moscovitch et al., 2016) suggest that repeated reactivation leads context-specific memories to become more gist-like. Reactivation may involve hippocampal-cortical interactions that offload memory traces to the cortex, thereby decreasing reliance on the hippocampus. Supporting this framework, Hindy et al. (2026) examined the reactivation of word lists while participants completed a distractor task and found that reactivation in the hippocampus, especially the anterior hippocampus, was associated with increased false recognition of semantic lures and decreased source-memory accuracy. They interpreted these findings as evidence that reactivation shifts memories towards gist over episodic detail, rather than improving memory overall.

In the present study, reactivation was not reliably correlated with memory during the same-day testing session. One week later, participants completed another session in which they watched and recalled another episode of the same series and then recalled the episode that they had viewed during the prior scan session. An exploratory analysis showed that reactivation during Session 1 was correlated with more gist-like memory one week later, with participants who showed stronger reactivation recalling more gist units compared to detail units from memory. This could reflect a purely time-dependent effect of reactivation, or an interaction of time with the intervening episode or the prior week’s recall. Future work should replicate this finding and disentangle these potential factors.

Consistent with these ideas, recent findings in both rodents and humans suggest that reactivation during wake is not a veridical playback of experiences to strengthen specific memories (Foster, 2017), but instead serves as a compositional computation to reorganize or integrate information and form new knowledge (Kurth-Nelson et al., 2023; Van Der Meer and Bendor, 2025). For instance, Liu et al. (2019) found that reactivation during awake rest can reorganize experiences based on how they have to be remembered for a memory test rather than in the order in which items were presented, suggesting a role for reactivation in transforming memory traces. Schlichting and Preston (2014) found that reactivation of learned face-object associations (AB pairs) predicted better subsequent learning of related content (BC associations and AC inferences). In addition, replay sequences in rodents seem to reflect all possible paths in an environment, potentially integrating recent experiences with alternative potential outcomes to form generalized representations of the space (Gupta et al., 2010; Stella et al., 2019; Widloski and Foster, 2022). These findings are consistent with prior work on memory reactivation during sleep, which similarly suggests that sleep supports abstraction (Walker, 2009; Santamaria et al., 2024; Zhao et al., 2024; Solomon et al., 2026).

This work also connects to behavioral research in humans showing that following the encoding of a narrative, people have a tendency to spontaneously think of thematically-related topics (Bellana et al., 2022). This suggests that reactivation of complex events may be directed towards the integration of new experiences with prior knowledge (McClelland and O’Reilly, 1995; Moscovitch et al., 2016). Thus, our finding that reactivation may be associated with more gist-like memory could reflect the reactivation of thematically-related information that is being integrated into the experience, or the integration of various events to form a more gist-like narrative representation. Future studies could build on these findings by examining whether post-learning reactivation modulates the real-time integration of prior knowledge during new narrative experiences, and whether integration in turn alters memory for previously learned events.

Finally, the heterogeneity of findings in the literature suggests that it may be overly simplistic to assume a uniform role of memory reactivation across all situations. For instance, reactivation during sleep and waking states may have different effects (Oudiette et al., 2013; Schapiro et al., 2018), and reactivation driven by reinforcements (Singer and Frank, 2009; Gruber et al., 2016) might also differ from spontaneous reactivation. Moreover, post-learning changes in functional interactions between brain regions (e.g., Tambini et al., 2010; Collins and Dickerson, 2019; Folvik et al., 2023) have been shown to correlate with memory performance, but these changes do not necessarily correspond to reactivation of experience-related neural representations. Further work is needed to characterize the differential effects of reactivation on memory across these contexts. With this study we begin to bridge the gap between separate accounts of offline reactivation and extend our understanding to the transformation of continuous narrative experiences.

## Acknowledgements

This research was supported by the National Institute on Aging Grant 1R03AG063224, the National Institute on Aging Training Grant Program in Aging (T32) AG000030-47, the Office of Naval Research (ONR) Grant N00014-17-1-2961, and the National Institute of Mental Health (NIMH) Training Grant Program (T32) MH112507.

